# Novel IRE1-dependent proinflammatory signaling controls tumor infiltration by myeloid cells

**DOI:** 10.1101/533018

**Authors:** Joanna Obacz, Jérôme Archambeau, Daria Sicari, Pierre Jean Le Reste, Raphael Pineau, Sophie Martin, Kim Barroso, Efstathios Vlachavas, Konstantinos Voutetakis, Tanya Fainsod-Levi, Akram Obiedat, Zvi Granot, Boaz Tirosh, Juhi Samal, Abhay Pandit, John Patterson, Qinping Zheng, Luc Négroni, Aristotelis Chatziioannou, Véronique Quillien, Tony Avril, Eric Chevet

**Affiliations:** Inserm U1242, University of Rennes, Rennes, France; Centre de lutte contre le cancer Eugène Marquis, Rennes, France; Neurosurgery dept, University Hospital of Rennes, 35000 Rennes, France; Institut de Génétique et de Biologie Moléculaire et Cellulaire, 67404 Illkirch, France; Centre National de la Recherche Scientifique, UMR7104, 67404 Illkirch, France; Institut National de la Santé et de la Recherche Médicale, U1258, 67404 Illkirch, France; Université de Strasbourg, 67404 Illkirch, France; e-NIOS PC, Kallithea-Athens, Greece; Department of Developmental Biology and Cancer Research, Institute for Medical Research Israel-Canada, Hebrew University Medical School, Jerusalem, Israel; Institute for Drug Research, School of Pharmacy, Faculty of Medicine, Hebrew University of Jerusalem, Jerusalem, Israel; Institute of Chemical Biology, NHRF, Athens, Greece; Department of Biochemistry and Biotechnology, University of Thessaly, Larissa, Greece; Department of Molecular Biology and Genetics, Democritus University of Thrace, 68100 Dragana, Greece; CÚRAM, Centre for Research in Medical Devices, National University of Ireland, Galway, Ireland; Fosun Orinove PharmaTech Inc., 3537 Old Conejo Road, Suite 104, Newbury Park, CA 91320, USA; Fosun Orinove PharmaTech Inc., Suite 211, Building A4, 218 Xinghu St., Suzhou Industrial Park, Jiangsu 215123, China; Rennes Brain Cancer Team (REACT), 35000 Rennes, France

**Author notes:** Correspondance to: Véronique Quillien, Tony Avril or Eric Chevet - INSERM U1242, University of Rennes, Centre de Lutte Contre le Cancer Eugene Marquis, Avenue de la bataille Flandres Dunkerque, 35042 Rennes, France.

**Keywords:** Unfolded Protein Response, proteostasis, inflammation

## Abstract

Tumor cells are exposed to intrinsic and environmental challenges that trigger endoplasmic reticulum (ER) homeostasis alteration, in turn leading to ER stress. To cope with this, tumor cells engage an adaptive signaling pathway, the unfolded protein response (UPR) thus promoting the acquisition of malignant features. As such, glioblastoma multiforme (GBM), the most aggressive primary brain tumors, exhibit constitutive UPR signals to sustain growth. Herein, we showed that signaling elicited by one of the UPR sensors, IRE1, promotes GBM tumor invasion, angiogenesis and infiltration by macrophages. Hence, high IRE1 activity in tumors predicts worse outcome. We further dissect IRE1-dependent mechanisms that shape the brain tumor immune microenvironment towards myeloid cells. We identify an IRE1-dependent signaling pathway that directly controls the expression/release of proinflammatory chemokines (CXCL2, IL6, IL8) leading to tumor cell-mediated chemoattraction of neutrophils and macrophages. This pathway requires XBP1 non-conventional mRNA splicing and XBP1s-dependent expression of the E2 ubiquitin enzyme UBE2D3. The latter contributes to the degradation of the NFκB inhibitor IκB, leading to the up-regulation of proinflammatory chemokines. Our work identifies a novel IRE1/UBE2D3 proinflammatory signaling axis instrumental to pro-tumoral immune regulation of GBM.

## INTRODUCTION

Perturbation of endoplasmic reticulum (ER) protein homeostasis (also known as proteostasis) is one of the hallmarks of highly proliferative or secretory cells. Moreover, many intrinsic and environmental challenges, such as low oxygen levels, pH decrease or nutrient shortage may also increase the risk of ER proteostasis imbalance, thus yielding ER stress. To cope with this, cells engage an adaptive response named the unfolded protein response (UPR) which aims at alleviating ER stress or inducing apoptosis when stress cannot be resolved (Chen and Brandizzi, 2013; Chevet et al., 2015; Dufey et al., 2014; Maurel et al., 2015). IRE1 is the most conserved UPR transducer, and harbors both serine/threonine kinase and endoribonuclease (RNase) activities. Upon ER stress, IRE dimerizes/oligomerizes leading to its trans-autophosphorylation and the subsequent activation of the RNase activity. IRE1 RNase catalyzes XBP1 mRNA non-conventional splicing (XBP1s) (Calfon et al., 2002) together with the tRNA ligase RTCB (Almanza et al., 2018) and degrades RNAs though the Regulated IRE1 dependent decay (RIDD) of RNA (Hollien and Weissman, 2006; Maurel et al., 2014). XBP1s is a potent transcription factor of genes involved in protein glycosylation, ER-associated degradation, protein folding, and lipid synthesis (Almanza et al., 2018; Hetz et al., 2011), while RIDD output controls cell fate under ER stress conditions (Maurel et al., 2014). Features of ER stress and UPR hyper-activation have been observed in many pathological situations including cancer. As such, the UPR has emerged as an adaptive mechanism supporting tumor progression and resistance to treatment by impacting almost all cancer hallmarks (Urra et al., 2016). Mounting evidence also suggests that UPR shapes tumor microenvironment by regulating angiogenesis, inflammation and host immune response (Logue et al., 2018; Obacz et al., 2017b).

The consequences of UPR signaling have been studied in various cancers such as breast, liver, lung, prostate, pancreas, and in glioblastoma multiforme (GBM), one of the most lethal and aggressive primary brain tumor with a median survival of 15-18 months despite invasive multimodal treatment (Le Reste et al., 2016). IRE1 contributes to GBM development by regulating tumor growth, migration, invasion and vascularization, partially through RIDD-mediated cleavage of SPARC mRNA (Auf et al., 2010; Dejeans et al., 2012). Loss of functional IRE1 signaling is also associated with decreased expression of proangiogenic and proinflammatory VEGFA, IL1β, IL6, and IL8 in GBM cells (Auf et al., 2010). Importantly, we have recently shown the pivotal role of IRE1 in the immune remodeling of GBM stroma, which engaged the antagonistic roles of XBP1s and RIDD (Lhomond et al., 2018). However, the precise IRE1-dependent mechanisms underlying the pro-tumoral immune response in GBM remain elusive.

Existing relationships between IRE1 signaling and expression of proinflammatory chemokines have been established in different cellular models. Indeed IRE1 signaling has been shown to promote the expression of proinflammatory chemokines through both XBP1s and JNK-dependent pathways (Martinon et al., 2010; Shanware et al., 2014) or through IRE1-mediated induction of GSK3β (Kim et al., 2015). IRE1 was also reported to interact with TRAF2, recruiting IκB kinase (IKK), which triggers in turn the phosphorylation and degradation of IκB, enabling the NFκB nuclear translocation (Hu et al., 2006). In addition, the IRE1-TRAF2 complex was shown to induce JNK phosphorylation and subsequent upregulation of proinflammatory genes through activator protein 1 (AP1) (Grootjans et al., 2016; Urano et al., 2000). Given that brain malignancies are infiltrated by a large number of immune cells, which modulate GBM aggressiveness and response to treatment, we aimed at investigating the molecular mechanisms by which IRE1 controls GBM immune infiltrate. We showed that IRE1 orchestrates the reciprocal communication between cancer and non-tumoral cells of the microenvironment by promoting the recruitment of myeloid cells to GBM. This involves a novel IRE1/UBE2D3 signaling mechanism that depends on XBP1 mRNA non-conventional splicing and regulates NFκB activation. This subsequently leads to the production of proinflammatory chemokines and the recruitment of immune/inflammatory cells to the tumor site.

## RESULTS

### Myeloid cells are recruited to GBM in an IRE1-dependent manner *in vitro* and *in vivo*

We previously demonstrated that IRE1 activity in GBM cells controlled the recruitment of inflammatory cells such as macrophages and microglial cells (Lhomond et al., 2018). Here, we sought to test whether this was also true for other immune cells including neutrophils and T cells from the myeloid and lymphoid lineages, respectively. To this end, we first interrogated whether the immune gene signatures characterizing microglia/macrophages (MM), polynuclear neutrophils (PN) or T cells (T) were associated with IRE1 activity as defined with our recently established categorization of tumors according to IRE1 signature (Lhomond et al., 2018) using the TCGA GBM transcriptome data (**Fig.1A, S1A and S1B**). We found that myeloid MM and PN gene signatures were strongly linked to the high IRE1 activity gene signature. Using flow cytometry, we confirmed the presence of both myeloid cell types in freshly dissociated samples from an in-house GBM cohort (n=65) by quantifying the expression of specific markers for each cell types (**Fig.1B**). MM constituted a majority of GBM infiltrates as described in (Badie and Schartner, 2000), while PN were found in approximately 20% of analyzed specimens, whereas T cell infiltration was rare (around 10%) (**Fig.1B**). This suggested that IRE1 signaling could contribute to attracting myeloid cells that in turn may promote tumor aggressiveness. To further test this hypothesis, monocytes and neutrophils attraction was assayed using Boyden chamber assays. To this end, myeloid cells including monocytes (Mo) and PN were isolated from the blood of healthy donors and PN were characterized based on their morphology and on the expression of CD14, CD15, CD66b and CD16 cell surface receptors (**Fig.S1C**). We also generated a GBM primary cell line RADH87 with stable overexpression of wild-type (WT) IRE1 or of a truncated IRE1 variant, Q780* (**Fig.S1D**). The latter mutation was shown to abrogate the IRE1 signaling and its downstream output towards XBP1 mRNA splicing, therefore resembling the characteristics of U87 DN cells (**Fig.S1D** (Lhomond et al., 2018)). As shown in **Fig.1C and 1D**, tumor cell conditioned media (TCCM) derived from U87 DN and RADH87 Q780* cells did not recapitulate the migratory abilities of Mo and PN in Boyden chamber assay, when compared to TCCM from parental U87 and RADH87 cells or RADH87 IRE1 WT cells. Lastly, we validated the role of IRE1 activity in myeloid cell recruitment *in vivo*. As such, mice were first injected with GL261 GBM cells. Fourteen days after injection, mice underwent surgical removal of their tumor, followed by insertion of either empty gel implant (referred to as “plug”) or plug containing the IRE1 inhibitor, MKC8866 (Logue et al., 2018; Reste et al., 2019). We showed that pharmacological inhibition of IRE1 activity with MKC8866 significantly decreased PN but not MM infiltration to the recurring GBM (**Fig.1E-F**). Overall, our results demonstrate that GBM are infiltrated by myeloid cells and this is mediated at least in part by IRE1 signaling.

**Figure 1.**
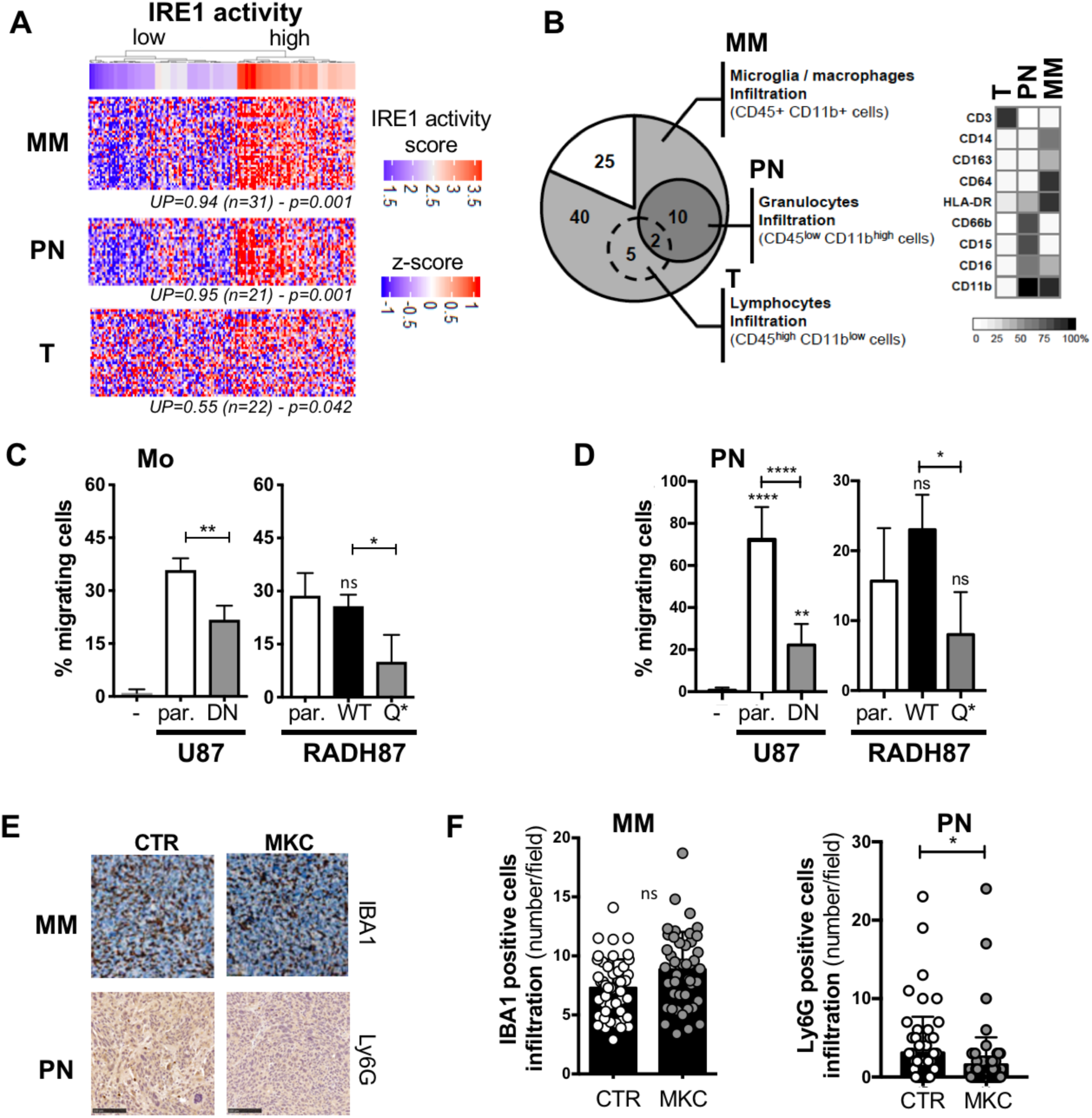
Impact of IRE1 on myeloid recruitment to GBM *in vitro* and *in vivo*. **A)** Hierarchical clustering of GBM patients (TCGA GBM cohort) based on high or low IRE1 activity obtained from (Lhomond et al., 2018) was confronted to immune cell markers for microglia/macrophage (MM), polynuclear neutrophils (PN) and T cells (T) derived from the literature. **B)** Total immune infiltrate of human GBM tissues (n=65) as analyzed by flow cytometry using anti-CD45 and anti-CD11b antibodies. Number of tumors infiltrated by specific leukocytes CD45+ populations are shown in circles. Deeper characterization of the different subtypes of immune CD45+ cells was performed combining the specific markers. **C-D)** Freshly isolated myeloid cells (i.e. monocytes (Mo) and polynuclear neutrophils (PN)) were placed in Boyden chambers towards TCCM from U87, U87 DN, parental RADH87 cells or RADH87 cells overexpressing IRE1 WT or Q780* mutant cells and incubated for 2 hours (for PN) or 16 hours (for Mo). Migrating cells were then quantified by flow cytometry using FCS/SSC parameters. Data are represented as percentage of myeloid cells migrated through the chamber compared to the initial number of cells placed in the insert (n=3, mean ± SD). (*): p<0.05, (**): p<0.01, (****): p<0.0001. **E)** Representative immunohistological analysis of myeloid infiltration in GBM specimens resected from mice treated with empty plug or plug with IRE1 inhibitor MKC8866. MM and PN were detected with anti-IBA1 and anti-Ly6G antibodies, respectively. Bar scale 100µm. **F)** Semi-quantitative analysis of IBA1 and Ly6G staining in tumors from (E). At least thirty random fields from tumor tissue and at least thirty random fields from tumor periphery were quantified for control (PLUG) and MKC-treated group. (*): p<0.05.

### IRE1 activity regulates the expression of myeloid-attracting chemokines

We next hypothesized that IRE1 activity in tumor cells could control the expression of specific chemokines, thus resulting in the attraction of myeloid cells. A number of cytokines/chemokines stimulating myeloid attraction was described in different cancer models, including CXCL1, CXCL2, CXCL5, CXCL7, IL6, and IL8 for PN (Powell and Huttenlocher, 2016) and CCL2, CCL3, CCL5, CCL8, IL6 and IL8 for MM (Guedes et al., 2018; Turner et al., 2014). Using transcriptome data from the TCGA GBM cohort, we demonstrated that tumors exhibiting high IRE1 activity express high mRNA levels of the aforementioned cytokines/chemokines (**Fig.2A**). Higher mRNA expression of these cytokines/chemokines was also observed in tumors with high levels of MM and PN myeloid markers, namely CD14 and CD15/CD16 (**Fig.2B and S2A**). To test whether any of those chemokines could be responsible for promoting the myeloid recruitment to GBM, freshly isolated Mo and PN were exposed to TCCM derived from various GBM (both primary and standard) cell lines. We then analyzed chemokines expression in those TCCM using ELISA-based assay and found that robust chemoattraction of myeloid cells correlated with elevated levels of CCL2, CXCL2, IL6 and IL8 (**Fig.2C**). As expected, PN (but not Mo) attraction was partially blocked by SB225002, a CXCR2 antagonist (**Fig.2D**). In line with our previous work (Auf et al., 2010), we showed that expression of myeloid attracting chemokines depended on IRE1 activity. The expression of CXCL2, IL6 and IL8 but not CCL2 mRNA was dramatically reduced in U87 DN and RADH87 Q780* cells when compared to control or WT-overexpressing cells (**Fig.2E**). Importantly, CXCL2, IL6 and IL8 mRNA levels were higher in GBM cell lines exhibiting high XBP1s activity (**Fig.2F**). These findings indicate that IRE1 signaling, possibly through XBP1s, is involved in the regulation of chemokines expression, which once secreted could be involved in myeloid recruitment to GBM tumors.

**Figure 2.**
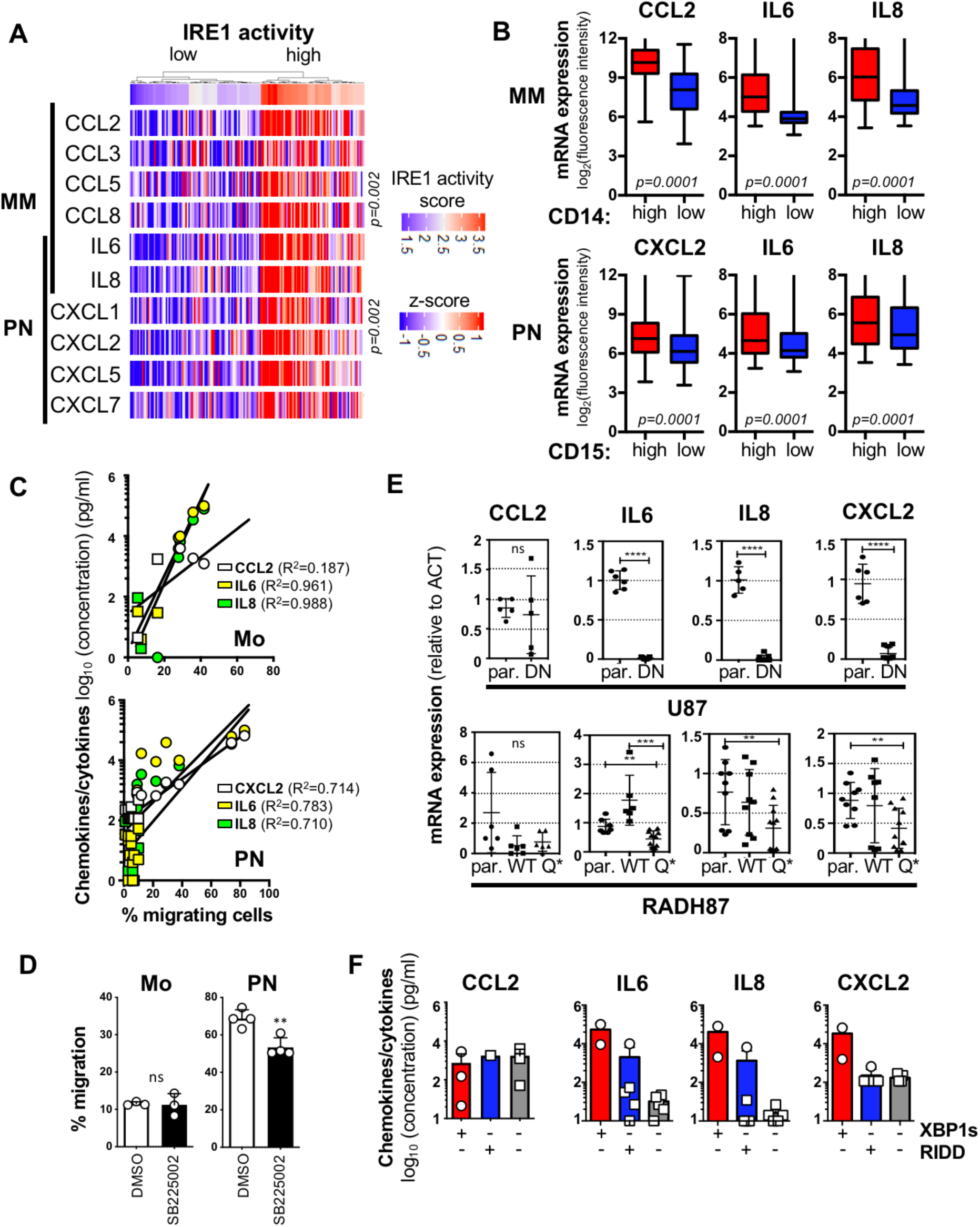
IRE1-mediated synthesis of myeloid-attracting chemokines. **A)** Hierarchical clustering of GBM patients (TCGA GBM cohort) based on high or low IRE1 activity confronted to the expression of cytokines/chemokines involved in myeloid cell chemoattraction. **B)** mRNA expression of CCL2, CXCL2, IL6 and IL8 in the population of tumors with high (red) or low (blue) MM and PN infiltration, as determined according to CD14 or CD15 levels, respectively. **C)** Correlation between CCL2, CXCL2, IL6 and IL8 secretion and Mo / PN migration towards TCCM from different GBM cell lines. **D)** Myeloid cell migration assay was performed as described in Fig.1C in the presence of DMSO or SB225002, a CXCR2 antagonist. **E)** Quantification of the expression of myeloid-attracting chemokines using RT-qPCR in U87, U87 DN, parental RADH87 or RADH87 cells overexpressing wild-type (WT) or Q780* IRE1 mutant. Data are representative of three independent experiments. (**): p<0.01, (***): p<0.001, (****): p<0.0001. **F)** Protein expression of CCL2, CXCL2, IL6 and IL8 in different GBM cell lines classified according to XBP1s or RIDD activity obtained from (Lhomond et al., 2018).

### UBE2D3 is a novel component of IRE1/XBP1s signaling involved in the regulation of myeloid infiltration in GBM through the activation of NF*κ*B

To test the involvement of XBP1s in chemokine production and since previous studies demonstrated that expression of these proinflammatory mediators is controlled by NFκB signaling (Kunsch and Rosen, 1993; Libermann and Baltimore, 1990), we used U87 DN cells in which we reintroduced XBP1s and evaluated the activation of NFκB through its phosphorylation and expression levels as well as through the expression of its inhibitor IκB. This revealed that overexpression of XBP1s in cells deficient for IRE1 signaling reduced the expression of IκB and increased NFκB phosphorylation, indicative of NFκB activation (**Fig.3A**). We then postulated that XBP1s could regulate molecular actors of the ubiquitin pathway controlling IκB degradation. To test this, we intersected known XBP1s target genes identified by ChIPseq (Chen et al., 2014) with all genes related to the ubiquitin system. We identified 13 candidates (**Fig.3B**) among which 7 were tightly interconnected (**Fig.3C**). Using transcriptome of the TCGA GBM cohort, we confirmed that mRNA expression of 3 of those genes, namely the E3 ubiquitin-protein ligase SYNV1, and the E2 ubiquitin-protein ligases UBE2D3 and UBE2J1 were positively correlated with XBP1 (**Fig.3D, S2B**). Moreover, UBE2D3 was previously described to be directly involved in IκB degradation (Gonen et al., 1999; Yaron et al., 1998). Furthermore, using MatInspector (Cartharius et al., 2005) we identified multiple XBP1s potential binding sites within the UBE2D3 promoter (**FigS2C**). We next confirmed that UBE2D3 expression was significantly upregulated in tumors with high IRE1 and XBP1s activities (**Fig.3E**). To validate our transcriptome-based findings, we measured UBE2D3 mRNA levels in cells with impaired IRE1 signaling. UBE2D3 expression was down-regulated in U87 and RADH87 cells deficient for either IRE1 activity (**Fig.3F**) or XBP1s (**Fig.3G**), respectively. High levels of UBE2D3 mRNA expression were also correlated with XBP1s expression in GBM cell lines subjected to tunicamycin-induced ER stress (**Fig.S2D**). At last, UBE2D3 expression was up-regulated in U87 and RADH87 cells deficient for IRE1 activity when XBP1s was re-introduced (**Fig.3H, S2E**). This indicated that XBP1s acts as a transcriptional regulator of UBE2D3 expression. Importantly, overexpression of UBE2D3 in GBM cell lines (**Fig.S3A and S3B**) led to the degradation of IκB and concomitant NFκB activation, as manifested by increased phosphorylation of NFκB protein (**Fig.3I, S3C and** (Wu et al., 2010)). This effect was not potentiated under tunicamycin treatment, suggesting that overexpression of UBE2D3 may bypass IRE1/XBP1s activation towards proinflammatory signals (**Fig.S3C**).

**Figure 3.**
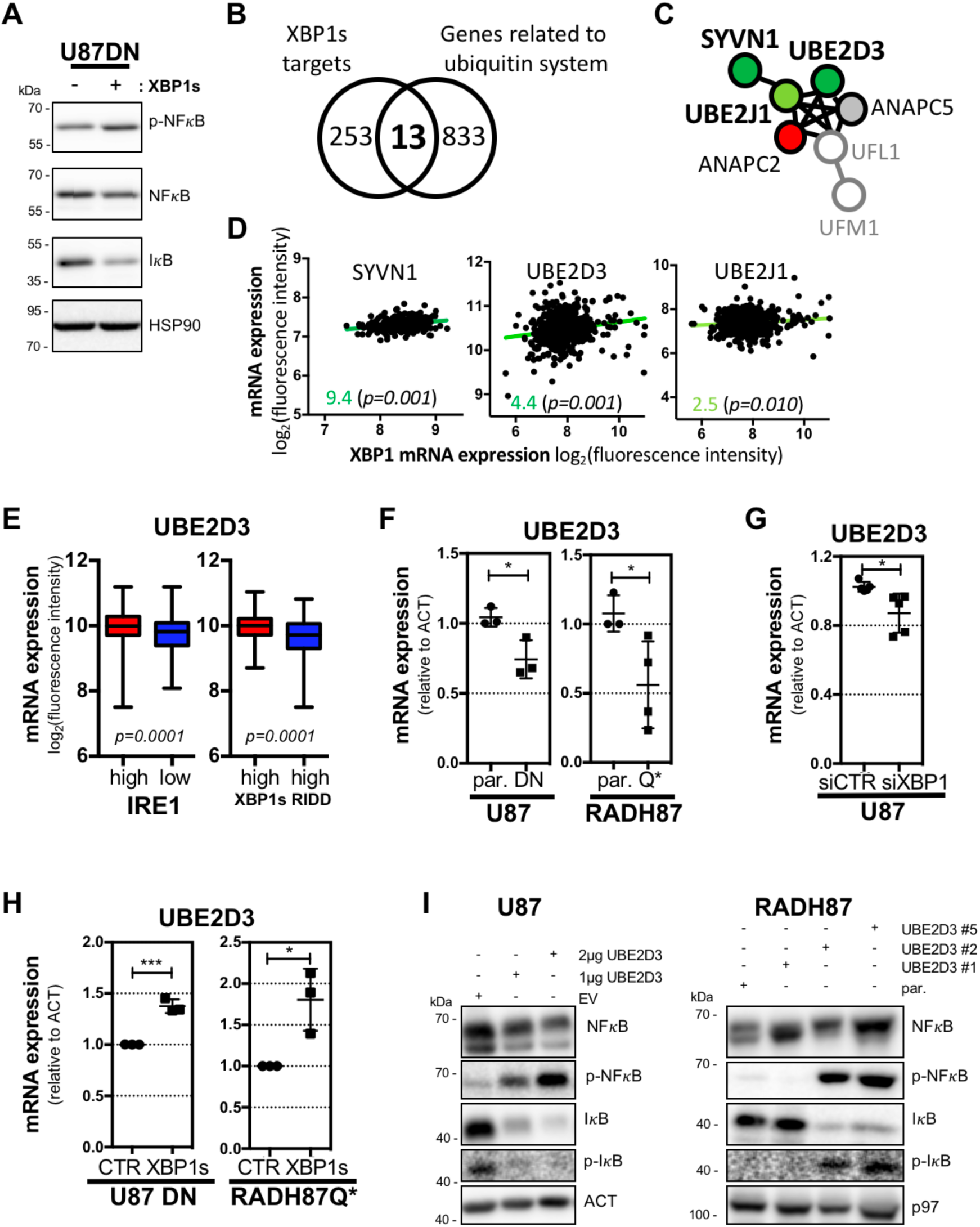
IRE1/XBP1-dependent regulation of UBE2D3 involved in NF*κ*B activation. **A)** Western blot analysis of NFκB and IκB expression as well as NFκB phosphorylation in U87 DN cells overexpressing XBP1s. HSP90 was used as control of protein loading. **B)** Venn diagram representation of the intersection of XBP1s target genes identified by chromatin immunoprecipitation sequencing approach described in (Chen et al., 2014) with genes related to ubiquitin system (GO term: 0016567). **C)** Functional protein-protein interaction network observed with 7 out of 13 genes determined using STRING database. Genes with positive or negative correlation with XBP1 expression were highlighted in green or red, respectively. **D)** Correlation of SYVN1, UBE2D3 or UBE2J1 mRNA expression with XBP1 mRNA expression in GBM specimens from the TCGA cohort. **E)** mRNA expression of UBE2D3 in GBM specimens from the TCGA cohort categorized according to their IRE1 activity, as in (Lhomond et al., 2018). **F)** Quantification of UBE2D3 mRNA expression by RT-qPCR in cells with active (U87 control (par) and RADH87 parental (par)) and inactive IRE1 signaling (U87 DN and RADH87 overexpressing Q780* mutant (IRE1_Q)). **G)** Quantitation of UBE2D3 expression using RT-qPCR in U87 cells silenced or not for XBP1. Data are representative of five independent experiments. (*): p<0.05. **H)** Quantitation of UBE2D3 expression with RT-qPCR in U87 DN cells and RADH87 cells overexpressing IRE1 Q780* mutant (RADH87 Q) and transfected with empty-vector (CTR) or XBP1s expression plasmid. Data are representative of three independent experiments. (*): p<0.05, (***): p<0.001. **I)** Western blot analysis of NFκB, phospho-NFκB, IκB and phospho-IκB in control (empty-vector, EV) and UBE2D3 overexpressing U87 and RADH87 cells. Actin (ACT) and VCP (p97) were used as loading control.

### UBE2D3 cooperates with the ubiquitin E3 ligase MIB1 to degrade IκB and trigger NFκB proinflammatory response

To identify the putative E3 ligase(s) involved in the degradation of IκB and/or activation of NFκB signaling as well as to investigate the global effect of UBE2D3 on proteins ubiquitination in GBM, we next carried out a label-free quantitative MS/MS analysis using cells with stable overexpression of UBE2D3 (**Fig.S3B**). Total proteins were extracted from RADH87 control and RADH87_UBE2D3 cells treated or not with tunicamycin. Precipitated proteins were then subjected to trypsin digestion, followed or not by purification of ubiquitin-derived diglycine (di-Gly) remnants and concomitant MS/MS analysis of both total and ubiquitinated peptides (**Fig.4A**). Among significantly up- and down-regulated proteins in UBE2D3 overexpressing cells compared to control, we identified a set of proteins involved in proteostasis control as well as in the inflammatory response (**Fig.4B**). Furthermore, we found a tight interaction between those ER-related entities that mainly function in protein metabolic processes (**Fig.4C**) and showed an enrichment in processes associated with membrane traffic (**Fig.S4A**), thus highlighting a key role of UBE2D3 in protein secretion.

**Figure 4.**
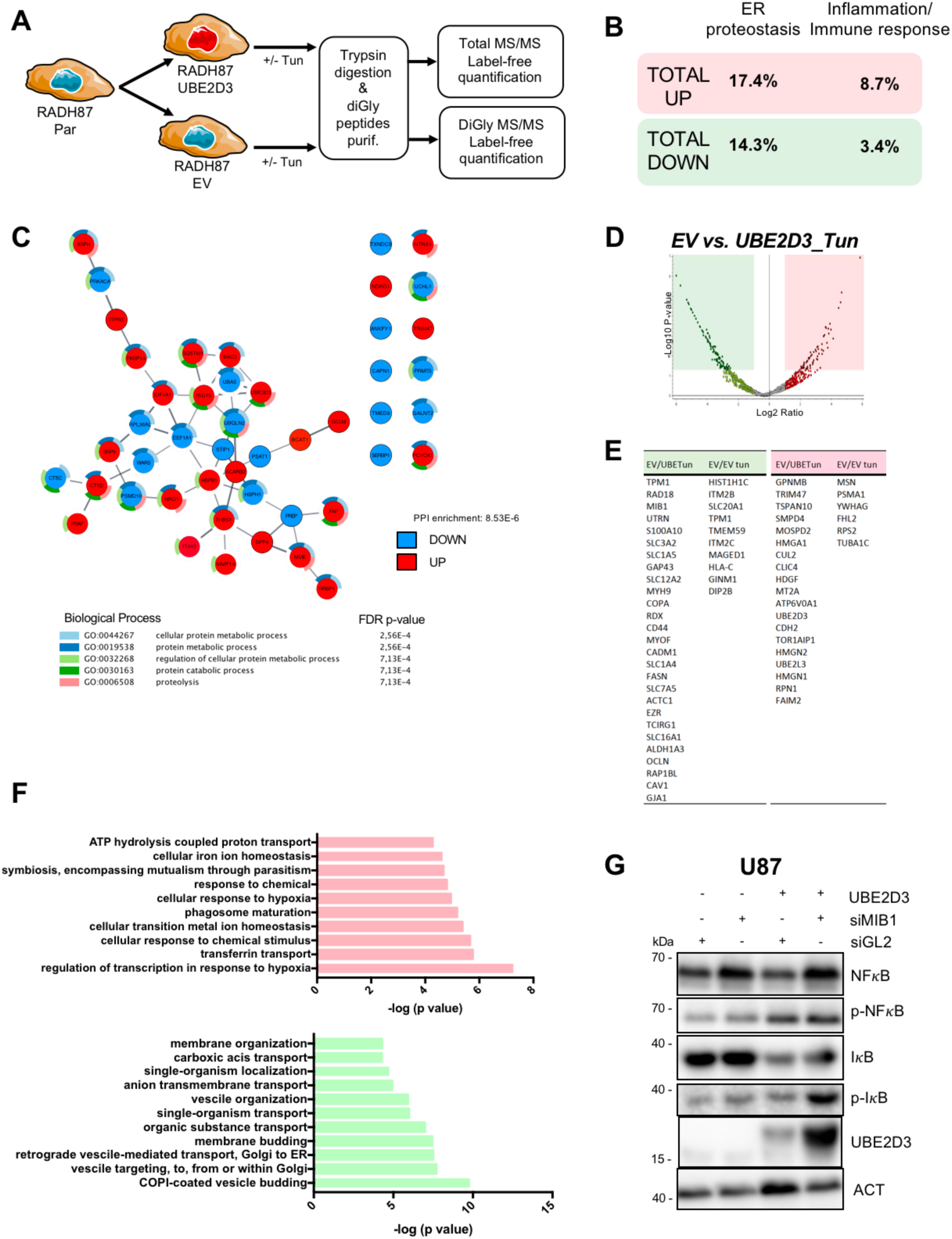
Impact of UBE2D3 on global proteins ubiquitination and its link to proteostasis. **A)** Schematic representation of the MS/MS experimental setup for the purification of ubiquitinated proteins from RADH87 control (EV) and UBE2D3 overexpressing cells. **B)** Percentage of up- and downregulated proteins related to ER proteostasis or inflammation/immune response in RADH87_UBE2D3 cells compared to RADH87 control. **C)** Representation of the ER-related protein network as identified in proteomics and list of statistically enriched GO biological processes. Indicated in colors are proteins, whose expression is modulated in UBE2D3 overexpressing cells (upregulated in red, downregulated in blue). **D)** Volcano plots of differentially ubiquitinated proteins purified from control (EV) or UBE2D3 overexpressing RADH87 cells exposed or not to ER stressor, tunicamycin (Tun). Proteins with p<0.05 and log ratio >2 or log ratio <-2 are delignated in pink and green, respectively. Dots represent purified peptides corresponding to the identified proteins. **E)** List of ubiquitinated proteins significantly upregulated (pink) or downregulated (green) in the indicated conditions. EV, control RADH87 cells; UBETun, RADH87_UBE2D3 cells treated with tunicamycin. **F)** Overrepresented (pink) and underrepresented (green) GO biological processes for the set of purified ubiquitinated proteins from RADH87_UBE2D3 cells exposed to tunicamycin (Tun) compared to RADH87 control (EV) cells. Fold enrichment values are represented as the minus base 10 log of their corresponding p values. GO-term enrichments analysis was performed using STRING database (https://string-db.org/). **G)** Western blot analysis of NFκB, phospho-NFκB, IκB, phospho-IκB and UBE2D3 in control (empty-vector, EV) and UBE2D3 overexpressing U87 cells after MIB1 down-regulation with a siRNA approach. Actin (ACT) was used as loading control.

We next analyzed peptides containing di-Gly ubiquitin remnants for each sample and compared the differentially ubiquitinated proteins between the most extreme conditions (RADH87 control (EV) cells cultivated without stress vs. RADH87_UBE2D3 cells treated with tunicamycin), reasoning that this would encompass the effect of UBE2D3 on protein ubiquitination in both basal and ER stress conditions (**Fig.4D**). We identified forty-five proteins, whose ubiquitination was significantly altered in the context of UBE2D3 overexpression and ER stress in GBM (**Fig.4E**). Functional enrichment analysis further revealed that UBE2D3 overexpression mainly mediated the ubiquitination of proteins involved in cellular response to environmental stress (chemicals or hypoxia), while it reduced the ubiquitination of proteins involved in membrane transport (in particular retrograde trafficking; **Fig.4F**), which further supports our findings on the role of UBE2D3 in the regulation of ER homeostasis and secretory pathway. Within the proteins purified from UBE2D3 overexpressing cells that showed divergent ubiquitination patterns compared to parental cells, we identified the E3 ligase MIB1. We thus focused our attention on that molecule since MIB1 was reported to participate in the control of NFκB activation (Liu et al., 2012) and also to interact with UBE2D3 E2 enzyme (van Wijk et al., 2009). Therefore, we investigated whether MIB1 cooperated with UBE2D3 to trigger the degradation of IκB, leading to the activation of NFκB signaling and subsequent synthesis of myeloid-attracting chemokines in GBM. siRNA-mediated MIB1 silencing was carried out in UBE2D3 overexpressing cells (**Fig.S4B**) and the effects on NFκB pathway activation monitored. We found that MIB1 silencing partially prevented the UBE2D3-mediated degradation of IκB protein (**Fig.4G**). Overall, these results suggest the existence of a signaling axis involving IRE1, XBP1s, UBE2D3 and MIB1 sufficient to trigger the activation of NFκB-dependent proinflammatory response in GBM.

### The IRE1/UBE2D3 axis controls myeloid recruitment to GBM through the activation of NF*κ*B proinflammatory response

To further document the role of UBE2D3 in myeloid-mediated immunity, we stratified the TCGA GBM cohort according to UBE2D3 mRNA expression levels and tested the expression of the main cytokines/chemokines stimulating myeloid attraction. Tumors with high UBE2D3 expression levels also expressed significantly higher levels of CXCL2, IL6 and IL8 mRNA (**Fig. 5A**). Next, we evaluated the expression of these myeloid-attracting chemokines using RT-qPCR in UBE2D3 overexpressing GBM cell lines. This showed that the expression of CXCL2, IL6 and IL8 was markedly upregulated upon UBE2D3 overexpression (**Fig.5B**). In addition, treatment of U87 cells with the NFκB inhibitor, JSH-23, which precludes the nuclear translocation of NFκB and consequently its transcriptional activity, impeded the UBE2D3-dependent increase in CXCL2, IL6 and IL8 mRNA expression (**Fig.5C**). We next found that MIB1 silencing partially prevented the UBE2D3-mediated upregulation of proinflammatory chemokines IL6 and IL8 in UBE2D3 overexpressing cells compared to control counterparts (**Fig.5D**). Finally, we confirmed that TCCM from UBE2D3 overexpressing cells prompted the increased Mo and PN migration in Boyden chamber assay compared to TCCM from control cells (**Fig.5E**). Therefore, based on the above findings, we demonstrated that the tightly regulated signaling circuit involving IRE1/XBP1s/UBE2B3/MIB1 controls NFκB-dependent chemokines synthesis and inflammatory response in GBM (**Fig.5F**).

**Figure 5.**
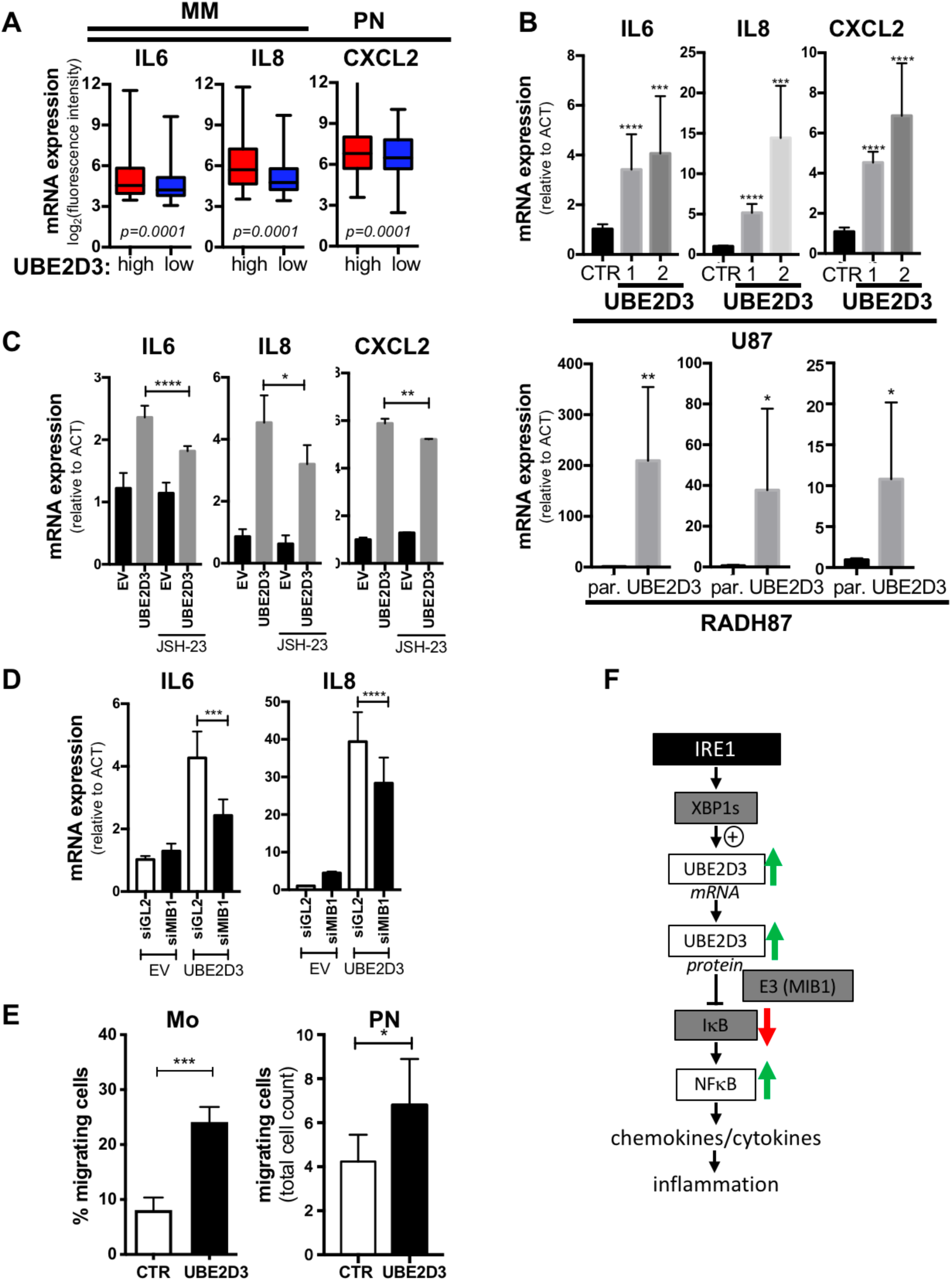
Impact of UBE2D3 on cytokines/chemokines synthesis. **A)** mRNA expression of IL6, IL8 and CXCL2 in GBM specimens from the TCGA cohort categorized according to their UBE2D3 expression (high: red and low: blue). **B)** Quantification of the expression of CXCL2, IL6 and IL8 using RT-qPCR in control (CTR) U87, parental (par.) RADH87, UBE2D3 overexpressing U87 and RADH87 cells. Data are representative of three independent experiments. (*): p<0.05, (**): p<0.01, (***): p<0.001, (****): p<0.0001 **C)** Quantification of the expression of CXCL2, IL6 and IL8 using RT-qPCR in U87 control (EV) or UBE2D3 overexpressing cells treated or not with 5µM NFκB pathway inhibitor, JSH-23. Data are representative of three independent experiments. (*): p<0.05, (**): p<0.01, (***): p<0.001, (****): p<0.0001. **D)** Quantification of IL6 and IL8 mRNA expression using RT-qPCR in U87 cells overexpressing UBE2D3 and silenced or not for MIB1. siGL2 was used as silencing control. (***): p<0.001, (****): p<0.0001. **E)** Migration of myeloid cells (Mo and PN) isolated from blood of healthy donors in Boyden chamber assay towards media conditioned by U87 control (ctl) and UBE2D3 overexpressing cells. Data are represented as total number of neutrophils migrated through the chamber (n=3, mean ± SD). (*): p<0.05. **F)** Schematic representation of the model describing the IRE1/UBE2D3 axis and its impact on inflammatory response in GBM. IRE1 through XBP1s controls UBE2D3 expression, that in turn together with E3 ligase of the ubiquitin system (including MIB1) directly degrades IκB, leading to the activation of NFκB signaling and its downstream targets.

### UBE2D3 controls pro-tumoral inflammation *in vivo* and increases GBM cell aggressiveness

To validate the importance of the IRE1/UBE2D3 signaling axis in GBM, we used a GBM syngeneic mouse model that allowed us to investigate the impact of UBE2D3 on myeloid infiltration to GBM *in vivo*. To this end, GL261 control and UBE2D3 overexpressing cells (denoted as GL261_UBE2D3 hereafter, **Fig.S5A**) were injected into the brain of immunocompetent C57BL/6 mice. Twenty-four days post-injections tumors were resected and subjected to the immunohistochemical analysis for quantifying the immune infiltrate. Interestingly, GL261_UBE2D3 cells produced larger tumors when compared to the control counterparts (**Fig.6A**), which was not attributed to the intrinsic characteristics of the cells, since both lines showed similar proliferation rate *in vitro* (**Fig.S5B**). This suggests that the growth advantage of UBE2D3 overexpressing tumors might emerge from the interaction with stroma and/or tumor microenvironment. In line with this observation, GL261_UBE2D3 tumors showed an elevated induction of NFκB expression (**Fig.6B**) and recruited significantly higher numbers of MM and PN to GBM tissue (**Fig.6C, 6D**), which confirmed our *in vitro* findings. Furthermore, using transcriptome data of two independent GBM cohorts, GBMmark and TCGA_GBM_LGG, we also demonstrated a strong correlation between UBE2D3 expression levels and the expression of a large number of proinflammatory cytokines/chemokines (**Fig.6E, S5C**). Intriguingly, high UBE2D3 expression was also associated with increased monocytes, T cells and M2-polarized macrophages infiltration, as determined by the expression levels of their specific surface receptors (**Fig.6F, S5D**). Next, to evaluate a clinical and prognostic relevance of UBE2D3 in GBM, we investigated the UBE2D3 expression in brain malignancies and showed that UBE2D3 is markedly increased in GBM specimens compared to low-grade gliomas (**Fig.6G**). We then compared the survival rates of patients whose tumors were characterized by high or low UBE2D3 expression and consequently found that higher expression of UBE2D3 is of poorer prognosis (**Fig.6G**). Lastly, we showed that temozolomide treatment had significantly less prominent effect on GL261_UBE2D3 cell death (**Fig.S5E**), which suggests that high UBE2D3 expression might render GBM cells more resistant to chemotherapy. Taken together, our findings unveil a novel IRE1-dependent mechanism promoting pro-tumoral inflammation, which integrates UPR signaling and ubiquitin system. Here, we demonstrated the existence of IRE1/UBE2D3 axis controlling the composition of GBM proinflammatory secretome through the activation of NFκB signaling (**Fig.6H**).

**Figure 6.**
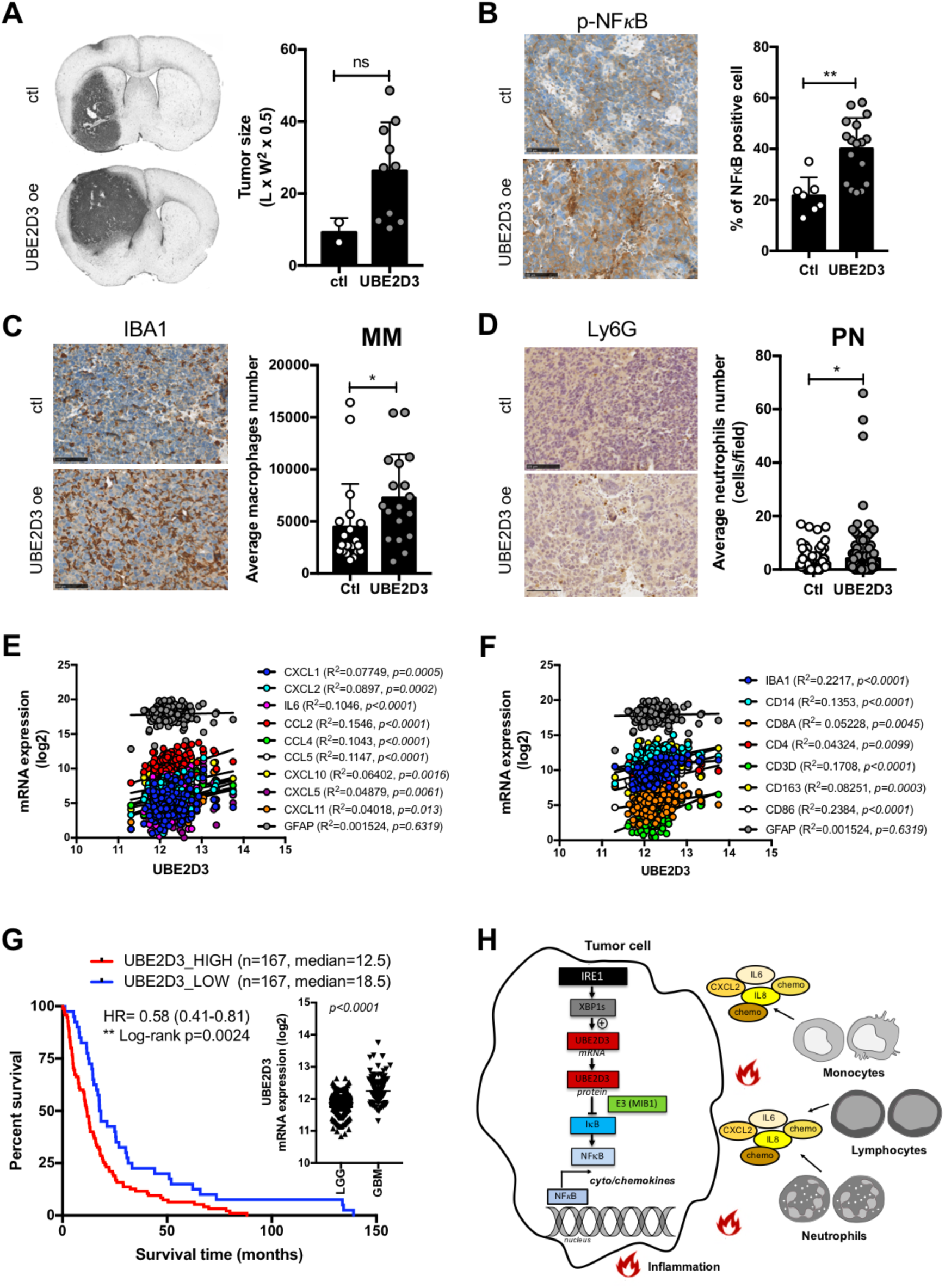
Impact of UBE2D3 overexpression on inflammation *in vivo*. **A)** Left panel: brain sections from mice injected with GL261 control (ctl) or GL261_UBE2D3 cells analyzed for vimentin expression by immunohistochemistry. Right panel: quantification of tumor volume between control (ctl) and UBE2D3 overexpressing group. **B-D)** Left panel: Representative immunohistological photographs of NFκB expression (B), macrophages/microglia infiltration (C) and neutrophils infiltration in GL261 control (ctl) or GL261_UBE2D3 tumors as detected by anti-NFκB, anti-IBA1 and anti-Ly6G antibodies, respectively. Right panel: semi-quantitative analysis of NFκB staining (B), IBA1 staining (C) and Ly6G staining (D) in GL261 control (ctl) and GL261_UBE2D3 tumors. At least forty random fields from tumor tissue and at least forty random fields from tumor periphery were quantified for control (ctl) and UBE2D3 overexpressing group. (*): p<0.05, (**): p<0.01. **E)** Correlation between UBE2D3 mRNA level and indicated cytokines/chemokines expression in GBM cohort. GFAP expression (astrocyte marker) was used as negative control. **F)** Correlation between UBE2D3 mRNA level and expression of indicated immune cell-specific receptors in GBM cohort i.e. CD3D, CD4 and CD8A for T cell markers; IBA1, CD14, CD86 and CD163 for MM markers. GFAP expression (astrocyte marker) was used as negative control. **G)** Comparison of UBE2D3 expression in low-grade gliomas (LGG) and glioblastoma (GBM) and its impact on patients’ survival. **H)** Schematic representation of IRE1/UBE2D3 axis in the regulation of pro-tumoral inflammation.

## DISCUSSION

Despite many advances in cancer research and the development of novel promising therapies, GBM still remains incurable. Given the systematic failure of the current therapeutic scheme, which involves maximal safe resection, followed by concomitant radiotherapy and chemotherapy (Stupp et al., 2005), there is an urgent need to better characterize the mechanisms underlying GBM development and progression, to identify novel therapeutic targets and design targeted innovative therapeutic approaches.

In the past years, we have characterized the relevance of UPR signaling, in particular IRE1 in GBM biology (Auf et al., 2010; Dejeans et al., 2012; Obacz et al., 2017a) and recently found that the characteristics of IRE1 signaling represented a predictive factor for GBM aggressiveness (Lhomond et al., 2018). As such, modulating ER stress signaling pathways poses an attractive therapeutic avenue for GBM treatment aiming at either increasing ER stress to levels that trigger apoptosis or decreasing adaptive signals (Obacz et al., 2017a). In addition to intrinsic aggressiveness of the GBM cells, the brain tumor microenvironment, that contains among others endothelial and immune cells, is emerging as a crucial regulator of brain cancer progression (Obacz et al., 2017a; Quail and Joyce, 2017). The most abundant immune cells in GBM microenvironment are tumor-associated macrophages and microglial cells that might reach up to 30% of the tumor mass and have been often linked to disease aggressiveness (Bingle et al., 2002; Hambardzumyan et al., 2016; Obacz et al., 2017b; Wei et al., 2013); however, brain tumors are also infiltrated by other immune cells such as myeloid dendritic cells (DCs), plasmacytoid DCs, T cells and neutrophils (Quail and Joyce, 2017).

In this work, we demonstrated that IRE1 signaling in tumor cells plays a key role in the regulation of GBM microenvironment, by promoting the recruitment of myeloid cells to the tumors. We previously found that IRE1 signaling was involved in the recruitment of macrophages and microglial cells to the tumors (Lhomond et al., 2018) and that IRE1 controlled the expression of proinflammatory chemokines (Logue et al., 2018; Pluquet et al., 2013). Herein, we showed that pharmacological inhibition of IRE1 signaling decreased the extend of PN infiltration to GBM *in vivo* (**Fig.1**). We also found that IRE1 activation in tumor cells correlated with higher expression of myeloid cells-attracting chemokines (**Fig.2**). To characterize IRE1-downstream signals responsible for this phenomenon, we showed that IRE1 tightly controlled the expression of UBE2D3 in GBM cells by engaging XBP1 mRNA non-conventional splicing (**Fig.3**). Then, we demonstrated that activation of the IRE1/XBP1s/UBE2D3 signaling axis was in part responsible for myeloid cell chemoattraction through the activation of proinflammatory NFκB (**Fig.5**). UBE2D3 is a ubiquitin-conjugating enzyme (E2) that together with ubiquitin-activating enzyme (E1) and ubiquitin ligase (E3) mediates the attachment of ubiquitin moieties to target proteins. This post-translational modification impacts on a broad range of biological processes, including protein quality control and trafficking, differentiation, cell division, signal transduction as well as inflammation (Glickman and Ciechanover, 2002; Mukhopadhyay and Riezman, 2007). UBE2D3 has been shown to control proteasomal degradation of among others p53 (Saville et al., 2004), cyclin D1 (Hattori et al., 2007), p12 subunit of DNA polymerase δ (Zhang et al., 2013) and IκBα (Wu et al., 2010; Yaron et al., 1998). It was also reported to mediate the ubiquitination of RIG-I, event required for its activation upon viral infection initiating the type I interferon (IFN)-dependent innate immune response (Shi et al., 2017). In this work, we further demonstrated a crucial role of UBE2D3 in the regulation of immunity/inflammation in pathological conditions, such as cancer. We found that UBE2D3 is overexpressed in GBM compared to low-grade gliomas and that its elevated expression correlated with high abundance of proinflammatory chemokines. We delineated a novel IRE1-dependent mechanism towards NFκB activation, which involves upregulation of UBE2D3 leading to the degradation of IκB, at least partially, through the E3 ubiquitin ligase MIB1, the subsequent nuclear translocation of NFκB and activation of its downstream signaling (**Fig.4**). Hence, IRE1 signaling controls the production of proinflammatory chemokines, including, as demonstrated here, CXCL2, IL6 and IL8. Once secreted, they not only sustain the pro-tumoral inflammatory microenvironment but can also mobilize the recruitment of immune cells to the tumor site further promoting cancer progression. As such, we showed *in vivo* that UBE2D3 overexpressing tumors were bigger in size and were infiltrated by significantly higher numbers of immune cells, such as MM and PN (**Fig.6**). However, our findings indicate that the aforementioned mechanism might be applicable to the infiltration by a large number of lymphocytes, highlighting the importance of understanding the IRE1/XBP1s/UBE2D3 axis in other cancer models, particularly in ‘immune hot’ tumors. Taken together, this study implies that targeting IRE1 signaling might impede glioblastoma aggressiveness by reducing tumor cell migration, invasion and angiogenesis (Auf et al., 2010; Lhomond et al., 2018), but also by neutralizing the pro-tumoral inflammation and immunity, hence opening a new avenue for therapeutic approaches for improving the efficacy of current immunotherapies.

## MATERIALS AND METHODS

### Antibodies and other reagents

All antibodies not specified below were purchased from BD Biosciences (Le Pont de Claix, France).The following primary antibodies were used: rabbit polyclonal anti-IRE1 (Santa Cruz), mouse monoclonal anti-XBP1s (Bio Legend), mouse monoclonal anti-KDEL (Enzo), mouse monoclonal anti-UBE2D3 (Abcam), rabbit monoclonal anti-NFκB p65 (Cell Signaling), rabbit monoclonal anti-phospho-NFκB p65 (Cell Signaling), rabbit polyclonal anti-IκB (Cell Signaling), rabbit monoclonal anti-phospho-IκB (Cell Signaling), rabbit monoclonal anti-actin (Sigma), mouse monoclonal anti-tubulin (Sigma) and mouse monoclonal anti-p97 (BD Transduction Laboratories). The secondary antibodies were horseradish peroxidase (HRP) conjugated polyclonal goat anti-rabbit IgG, HRP-conjugated polyclonal goat anti-mouse IgG and HRP-conjugated polyclonal rabbit anti-goat IgG (all Dako). Other reagents not specified below were purchased from Sigma-Aldrich (St Quentin Fallavier, France).

### Cell culture and treatments

The U87, U87 IRE1.NCK DN cells (referred to as U87 DN) (Drogat 2007), U251 and GL261 cell lines (all from ATCC) were grown in Dulbecco’s modified Eagle’s medium (DMED) (Invitrogen) supplemented with 10% fetal bovine serum (FBS) (Lonza) in a 5% CO_2_ humidified atmosphere at 37°C. Primary GBM cell lines were generated as previously described (Avril et al., 2012; Lhomond et al., 2018). For certain tumors, the two types of cultures have been established, adherent cell lines (RADH) grown in DMEM supplemented with 10% FCS and neurospheres (RNS, enriched in cancer stem cells) grown in DMEM/Ham’s F12 GlutaMAX (Life Technologies) supplemented with B27 and N2 additives (Invitrogen, Cergy Pontoise, France), EGF (20 ng/ml) and FGF2 (20 ng/ml) (Peprotech, Tebu-Bio) as described in (Avril et al., 2012). For transient overexpression or silencing, cells were transfected using Lipofectamine 2000 or Lipofectamine LTX (Thermo Fisher Scientific) for plasmids and Lipofectamine RNAiMAX Transfection Reagent (Thermo Fisher Scientific) for siRNA, according to the manual. To induce ER stress, cells were treated with 5 μg/mL Tunicamycin (Tun) (Calbiochem) for the indicated time periods. For NFκB pathway inhibition, 5µM JSH-23 (Sigma) was used for 16 hours.

### Patient samples and data

GBMmark cohort has been generated as previously described (Lhomond et al., 2018). Briefly, tumors were obtained from the processing of biological samples through the Centre de Ressources Biologiques (CRB) Santé of Rennes BB-0033-00056. The research protocol was conducted under French legal guidelines and fulfilled the requirements of the local institutional ethics committee. The quantification procedure of mRNA abundance is described in the “Microarray data analysis” section. Messenger RNA expression data were also assessed from the publicly available GBM dataset of The Cancer Genome Atlas (TCGA) (Consortium et al., 2007; consortium, 2008) from the NCBI website platform https://gdc-portal.nci.nih.gov/ and from the TCGA_GBM_LGG dataset obtained from GlioVis online tool (http://gliovis.bioinfo.cnio.es/).

### Microarray data analysis

Complete gene expression analysis of the GBMmark microarray Agilent dataset (GEO) was performed with R (R version 3.5.0)/Bioconductor software (Huber et al., 2015). Firstly, the raw data obtained from the public repository ArrayExpress (E-MTAB-6326) (Kolesnikov et al., 2015) were pre-processed (background correction and quantile normalization) using the limma R package (Ritchie et al., 2015). Next, non-expressed probesets in the majority of the samples (that is, probesets expressed in less than 10% of the total number of patients) were filtered out, in order to remove consistently non-expressed genes. To shed light on the molecular mechanisms involved in the IRE1-UBE2D3 signaling axis, the aforementioned list of DE genes, was used as an input to the BioInfoMiner interpretation web platform (Koutsandreas et al., 2016; Lhomond et al., 2018), which performs automated, network analysis of functional terms, integrating semantic information from different biomedical ontologies and controlled vocabularies such as Gene Ontology (GO), Reactome, Human Phenotype Ontology (HPO) and many others.

### Semi-quantitative PCR and Quantitative real-time PCR (qPCR)

Total RNA was extracted using the Trizol reagent (Invitrogen). All RNAs were reverse transcribed with Maxima Reverse Transcriptase (Thermo Scientific), according to manufacturer protocol. All PCR reactions were performed with a MJ Mini thermal cycler from Biorad (Hercules) and qPCR with a QuantStudio™ 5 Real-Time PCR Systems from Thermo Fisher Scientific and the PowerUp™ SYBR Green Master Mix (Thermo Fisher Scientific). Experiments were performed with at least triplicates for each data point. Each sample was normalized on the basis of its expression of the GAPDH or actin gene using 2^ΔΔCT^-method. The primers pairs used for this study are listed in **Table S1**.

### Mass spectrometry

RADH87 parental and RADH87_UBE2D3 cells were lysed with lysis buffer composed of 20 mM Tris pH 8, 1.5 mM EDTA, 150 mM NaCl, 1% Triton X-100, 0.1% SDS, 15µM MG132, 10mM NEM (N-ethylmaleimide), 10µM deubiquitinating enzymes inhibitors (DUBi, PR-619), supplemented with proteases and phosphatases inhibitor cocktails (Roche). Total proteins were precipitated overnight with 80% ice-cold acetone. Protein pellets were then washed 3 times with 80% acetone, followed by centrifugation at 500 rpm for 30 mins at 4°C. Samples were alkylated and digested with trypsin at 37°C overnight and ubiquitinated peptides were enriched with PTMScan Ubiquitin Remnant Motif (K-ε-GG) Kit (Cell Signaling Technology). After Sep Pak desalting, peptides were analyzed using an Ultimate 3000 nano-RSLC (Thermo Fisher Scientific) coupled in line with an Orbitrap ELITE (Thermo Scientific). Each sample was analyzed in triplicate. Briefly, peptides were separated on a C18 nano-column with a linear gradient of acetonitrile and analyzed in a Top 20 CID (Collision-induced dissociation) data-dependent mass spectrometry. Data were processed by database searching against Human Uniprot Proteome database using Proteome Discoverer 2.2 software (Thermo Fisher Scientific). Precursor and fragment mass tolerance were set at 7 ppm and 0.6 Da respectively. Trypsin was set as enzyme, and up to 2 missed cleavages were allowed. Oxidation (M, +15.995), GG (K, +114.043) were set as variable modification and Carbamidomethylation (C) as fixed modification. Proteins were filtered with False Discovery Rate <1% (high confidence). Lastly, quantitative values were obtained from Extracted Ion Chromatogram (XIC) and p-values were determined by ANOVA with Precursor Ions Quantifier node in Proteome Discoverer.

### Immunohistochemistry

Tumor tissues were fixed in 10% neutral buffered formalin, embedded in paraffin, cut into 4-μm thick sections, mounted on slides, deparaffinized in xylene and rehydrated into PBS through a graded ethanol series. Endogenous peroxidase activity was quenched in 3% hydrogen peroxide (Roche) in PBS for 15 minutes. The IHC labeling were carried out using the H2P2 imaging platform of the faculty of Rennes.

### Flow cytometry analyses

GBM specimens were dissociated using the gentle MACS dissociator (Miltenyi Biotec) according to manufacturer’s recommendations and cells were directly used for flow cytometry analysis. Cells were washed in PBS supplemented with 2% FBS and incubated with saturating concentrations of human immunoglobulins and fluorescent-labelled primary antibodies as indicated for 30 minutes at 4°C. Cells were then washed with PBS 2% FBS and analyzed by flow cytometry using a FACSCanto and Novocyte flow cytometer (BD Biosciences and Acea Biosciences). The population of interest was gated according to its FSC/SSC criteria. The dead cell population was excluded using 7AAD staining (BD Biosciences). Data were analyzed with the FACSDiva (BD Biosciences). GBM specimens with more than 2% stained cells of total viable cells were considered positive for the immune marker of interest.

### Myeloid chemoattraction assay

PBMC (source of monocytes) were isolated from peripheral blood of healthy donors with lymphocyte separation medium (Lonza) according to manufacturer’s recommendations. PN were isolated from peripheral blood of healthy donors using MACSXPRESS Neutrophil Isolation Kit (Miltenyi Biotec), according to manufacturer protocol. PBMC were used as source for Mo. Mo and PN were then washed in DMEM and placed in 3 μm Boyden chambers (Merck Millipore, France), at the concentration of 0.5 x 10^6^ cells/chamber in 200 µl DMEM. Chambers were simultaneously placed in 500 µl of either control DMEM or TCCM as indicated, and incubated for 2 hrs (for PN) and 16 hrs (for Mo) at 37°C. The migrated myeloid (under the Boyden chambers) were collected, washed in PBS and analyzed by flow cytometry using a Novocyte flow cytometer (Acea Biosciences). The population of interest was gated according to its FSC/SSC criteria. The relative number of migrated cells was estimated by flow cytometry by counting the number of cells per microliter. For CXCR2 inhibition with an antagonist SB225002, the drug was added and maintained during the migration assay.

### Syngeneic mouse model and inflammation

Eight-weeks old male C57BL/6 mice were housed in an animal care unit authorized by the French Ministries of Agriculture and Research (Biosit, Rennes, France - Agreement No. B35-238-40). The protocol used was as previously described (Auf et al., 2010). Cell implantations were at 2 mm lateral to the bregma and 3 mm in depth using GL261 (control) or GL261_UBE2D3 cells. Mice were daily clinically monitored and sacrificed twenty-four days post injection. Mouse brains were collected, fixed in 4% formaldehyde solution and paraffin embedded for histological analysis using anti-vimentin antibody (Interchim) to visualize the tumor masses. Tumor volume was then estimated by measuring the length (L) and width (W) of each tumor and was calculated using the following formula (L × W^2^ × 0.5). Immune cells infiltration was monitored by immunohistochemistry using rat anti-mouse Ly6G antibody (BD Biosciences) for neutrophils, anti-IBA1 (Wako) for macrophages/microglia, while NFκB level was determined with rabbit monoclonal anti-NFκB p65 antibody (Cell Signaling). Imaging was carried out using a Axioplan 2 epifluorescent microscope (Zeiss) equipped with a digital camera Axiocam (Zeiss).

### Statistical analyses

Graphs and statistical analyses were performed using GraphPad Prism 7.0 software (GraphPad Software). Data are presented as mean ± SD or SEM of at least three independent experiments. Statistical significance (p<0.05 or less) was determined using a paired or unpaired t-test or ANOVA as appropriate, while comparison of survival curves was done using log-rank (Mantel-Cox) test. Significant variations were represented by asterisks above the corresponding bar when comparing the test with the control condition or above the line when comparing the two indicated conditions.

All other methods can be found in Supplemental Material.

## Supporting information

Obacz supplemental

## Acknowledgements

We thank the Biosit histopathology H2P2 platform (Université de Rennes 1, France) and Florence Jouan for immunohistochemical analyses of tumor xenografts; the Biosit ARCHE animal facility (Université de Rennes 1) for animal housing. This work was funded by grants from INSERM, Institut National du Cancer (INCa), Région Bretagne, Rennes Métropole, Fondation pour la recherche Médicale (FRM; équipe labellisée 2018), EU H2020 MSCA ITN-675448 (TRAINERS), la Ligue Contre le Cancer Comités d’Ille-et-Vilaine, des Côtes d’Armor et du Morbihan and MSCA RISE-734749 (INSPIRED) to EC; PROMISE, 12CHN 204 Bilateral Greece-China Research Program of the Hellenic General Secretariat of Research and Technology and the Chinese Ministry of Research and Technology sponsored by the Program “Competitiveness and Entrepreneurship,” Priority Health of the Peripheral Entrepreneurial Program of Attiki to AC. JO was supported by a post-doctoral fellowship from Région Bretagne. DS was supported by an AIRC fellowship for Abroad.

## Notes

#### Summary of Updates

The whole manuscript has been completed to extend the study to both neutrophils and macrophages (myeloid cells).

