## Supplementary material for "Novel IRE1-dependent proinflammatory signaling controls tumor infiltration by myeloid cells": Obacz supplemental

#### MATERIALS AND METHODS

***Plasmids and siRNAs –*** pCMV3-UBE2D3-Flag expression plasmid was purchased from Sino Biological (Interchim, Montluçon, France), while pCMV5-Flag-XBP1s from Addgene (Teddington, UK). siRNA targeting the expression of MIB1 was obtained from Thermo Fisher Scientific.

***Chemokines secretion analysis –*** To evaluate the concentration of chemokines in tumor conditioned media, a Bio-Plex Multiplex immunoassays (Bio-Rad, Marnes-la-Coquette, France) was used following manufacturer protocol.

***Generation of stable cell lines –*** For stable overexpression of UBE2D3 protein, RADH87 and GL261 cells were transfected with 3μg of pCMV3-UBE2D3-Flag expression plasmid using Lipofectamine LTX and Lipofectamine 2000, respectively (Thermo Fisher Scientific), following the manufacturer protocol. RADH87 and GL261 cells stably overexpressing UBE2D3 (referred to as RADH87_UBE2D3 and GL261_UBE2D3 hereafter) were selected using 300μg/mL and 500μg/mL hygromycin B, respectively (Thermo Fisher Scientific). Monoclonal cell populations were obtained using limited dilution protocol after 10 days of antibiotic selection. Single-cells clones were expanded and UBE2D3 expression was analyzed by Western blot using anti-UBE2D3 (Abcam) or anti-Flag (Sigma) antibodies. Transfections of GBM adherent primary cell lines with IRE1 WT and Q780* constructs (Lhomond 2018) were performed using Lipofectamine LTX (Thermo Fisher Scientific), according to the manufacturer's instructions. IRE1 WT and Q780* overexpressing cells were selected using puromycin (Thermo Fisher Scientific).

***Cell proliferation*** – Parental and UBE2D3 overexpressing GL261 cells were cultured in 96-well plates at the concentration of $1x10^3$ cells/well and the growth rate was measured daily for

7 days. For the temozolomide (TMZ) sensitivity, cells were seeded at the concentration of $5x10^3$ cells/well and treated with the increasing concentrations of TMZ, ranging from 0-1000µM for 6 days. For both experiments, twenty microliters of the WST1 reagent (Roche) was added to the cells, followed by 4 hours incubation at 37°C and optic densities (OD) measurements with Tecan Infinite® 200 Pro spectrophotometry (Thermo Fisher Scientific) at 450 nm and 595 nm. Specific OD was obtained by calculating the difference between the OD at 450 nm and that at 595 nm and compared to the specific OD determined for DMEM alone.

#### FIGURE LEGEND

**Figure S1. Impact of IRE1 on myeloid recruitment to GBM *in vitro* and *in vivo*. A)** Barcode plot representation of the enrichment of immune cell gene signatures characterizing polynuclear neutrophils (PN), microglia/macrophages (MM) and T cells observed in GBM specimens (TCGA cohort) based on high or low IRE1 activity obtained from (Lhomond et al., 2018). **B)** mRNA expression of specific markers CD14, CD15 and CD4 corresponding to immune MM, PN and T cells in GBM specimens from the TCGA cohort categorized according to their IRE1 activity. **C)** Characterization of neutrophils isolated from blood using flow cytometry based on the expression of specific markers as designated. **D)** Western blot analysis of the expression of wild-type (WT) or Q780* IRE1, as well as that of XBP1s in RADH87 primary cell line exposed to tunicamycin treatment (Tun, 5 µg/mL for 6h). Actin (ACT) was used as loading control.

**Figure S2. Identification of neutrophil-attracting chemokines and IRE1/XBP1-dependent regulation of UBE2D3 in GBM. A)** mRNA expression of CCL3, CCL5, CCL8 (chemokines for MM) and CXCL1, CXCL5 and CXCL7 (chemokines for PN) in the population of tumors with high (red) or low (blue) MM and PN infiltration, as determined according to CD14 or CD16 levels respectively. **B)** Correlation of ANAPC2 and ANAPC5 mRNA expression with XBP1 mRNA expression in GBM specimens from the TCGA cohort. **C)** Graphical representation of exemplary transcription factors binding to the UBE2D3 promoter as analyzed with MatInspector. Putative XBP1 binding sites are delineated in blue (XBBF, X-box binding factors). **D)** Correlation between UBE2D3 and XBP1s mRNA expression in U87 (blue line) and RADH87 (green line) cells treated with 2.5 µg/ml tunicamycin for the time course of 1-24 hours. **E)** Correlation between UBE2D3 and XBP1s mRNA level in a panel of GBM cell lines transfected or not with plasmid coding for XBP1s as quantified by RT-qPCR.

**Figure S3. Modulation of UBE2D3 expression and its impact of NFκB signaling. A)** RT-qPCR validation of UBE2D3 gene overexpression in U87 cells. Cells were transfected with empty- vector (EV/CTR) or pCMV3-UBE2D3-Flag expression plasmid. The level of UBE2D3 mRNA was measured. **B)** Western blot validation of UBE2D3 overexpression in U87 and RAD87 cells. The level of UBE2D3 protein was measured with anti-UBE2D3 and anti-FLAG antibodies. **C)** Western blot analysis of NFκB, phospho-NFκB, IκB and phospoho-IκB in U87 and RADH87 control (empty-vector, EV) cells and cells with transient UBE2D3 overexpression under basal and ER stress conditions (Tun, tunicamycin; 6 hrs). Actin (ACT) was used as loading control.

**Figure S4. Impact of UBE2D3 on global proteins ubiquitination and its link to proteostasis. A)** Representation of the ER-related protein network as identified in proteomics and list of statistically enriched GO cellular components. Indicated in colors are proteins, whose expression is modulated in UBE2D3 overexpressing cells (upregulated in red, downregulated in blue). **B)** Quantification of MIB1 and UBE2D3 mRNA level with RT-qPCR in RADH87 control (empty-vector, EV) and UBE2D3 overexpressing cells after silencing for MIB1. (**): $p<0.01$, (***): $p<0.001$, (****): $p<0.0001$.

**Figure S5. Effect of UBE2D3 overexpression on GBM aggressiveness. A)** Western blot analysis of UBE2D3 protein overexpression in five different stable lines derived from GL261 parental cells transfected with pCMV3-UBE2D3-Flag expression plasmid. The level of UBE2D3 protein was measured with anti-Flag antibody. **B)** Proliferative rate of GL261 parental and UBE2D3 overexpressing cells as determined using WST1-based colorimetric assay. The proliferation index was quantified by calculating the difference between the absorbance at 450 nm and that at 595 nm. **C)** Correlation between UBE2D3 mRNA level and indicated cyto/chemokines expression in GBMmark cohort. GFAP expression (astrocyte marker) was used as negative control. **D)** Correlation between UBE2D3 mRNA level and expression of indicated immune cell-specific receptors in GBMmark cohort. GFAP expression (astrocyte marker) was used as negative control. **E)** The sensitivity of GL261 and GL261_UBE2D3 cells to the temozolomide treatment ranging from 0 to 1000µM, as quantified with WST1-based colorimetric assay. The proliferation index was quantified by calculating the difference between the absorbance at 450 nm and that at 595 nm and was normalized to DMSO treatment alone.

**Table S1. Primers used in the study**

| Gene | Forward primer | Reverse primer |
|---|---|---|
| **ACT** | 5'-CATGGGTGGAATCATAATGG-3' | 5- AGCACTGTGTTGCGCTACAG -3 |
| **CXCL2** | 5'-CTGCGCTGCCAGTGCTT-3' | 5'-CCTTCACACTTTGGATGTTCTTGA-3' |
| **GAPDH** | 5'-AAGGTGAAGGTCGGAGTCAA-3' | 5'-CATGGGTGGAATCATAATGG-3' |
| **IL6** | 5'-GGTACATCCTCGACGGCATCT-3' -3 | 5'-GTGCCTCTTTGCTGCTTTCAC-3' |
| **IL8** | 5'-TGGCAGCCTTCCTGATTTCT-3' | 5'-GGGTGGAAAGGTTTGGAGTATG-3' |
| **MIB1** | 5′-ACTGGCAGTGGGAAGATCAA-3′ | 5′-CATATGCTGCGCTATGTGGG-3′ |
| **IRE1** | 5'-GCCACCCTGCAAGAGTATGT-3' | 5'-ATGTTGAGGGAGTGGAGGTG-3' |
| **UBE2D3** | 5'-CCGGACCTTTGAGCATACAC-3' | 5'-GCCTTGATATGGGCTGTCAT-3' |
| **XBP1tot** | 5'-CCTGGTTCTCAACTACAAGGC-3' | 5'-AGTAGCAGCTCAGACTGCCA-3' |
| **XBP1s** | 5'-TGCTGAGTCCGCAGCAGGTG-3' | 5'-GCTGGCAGGCTCTGGGGAAG-3' |

### Figure S1

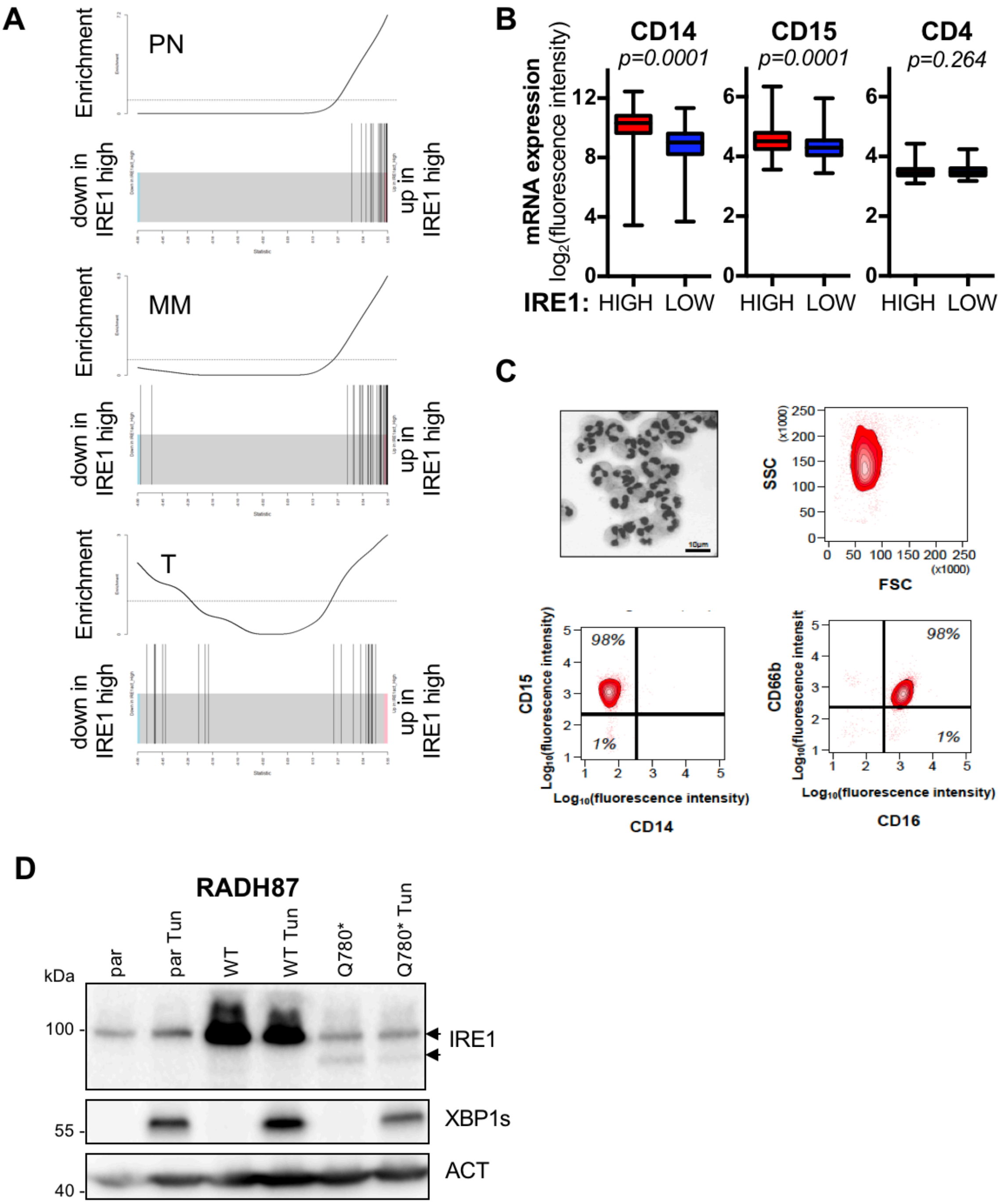

A
Enrichment
PN
down in IRE1 high
up in IRE1 high
MM
T
B
CD14
p=0.0001
CD15
p=0.0001
CD4
p=0.264
mRNA expression
log2(fluorescence intensity)
IRE1: HIGH LOW HIGH LOW HIGH LOW
C
SSC
FSC
(x1000)
10µm
CD15
98%
1%
Log10(fluorescence intensity)
CD14
CD66b
CD16
D
RADH87
par
par Tun
WT
WT Tun
Q780*
Q780* Tun
kDa
100
55
40
IRE1
XBP1s
ACT

#### Figure S2

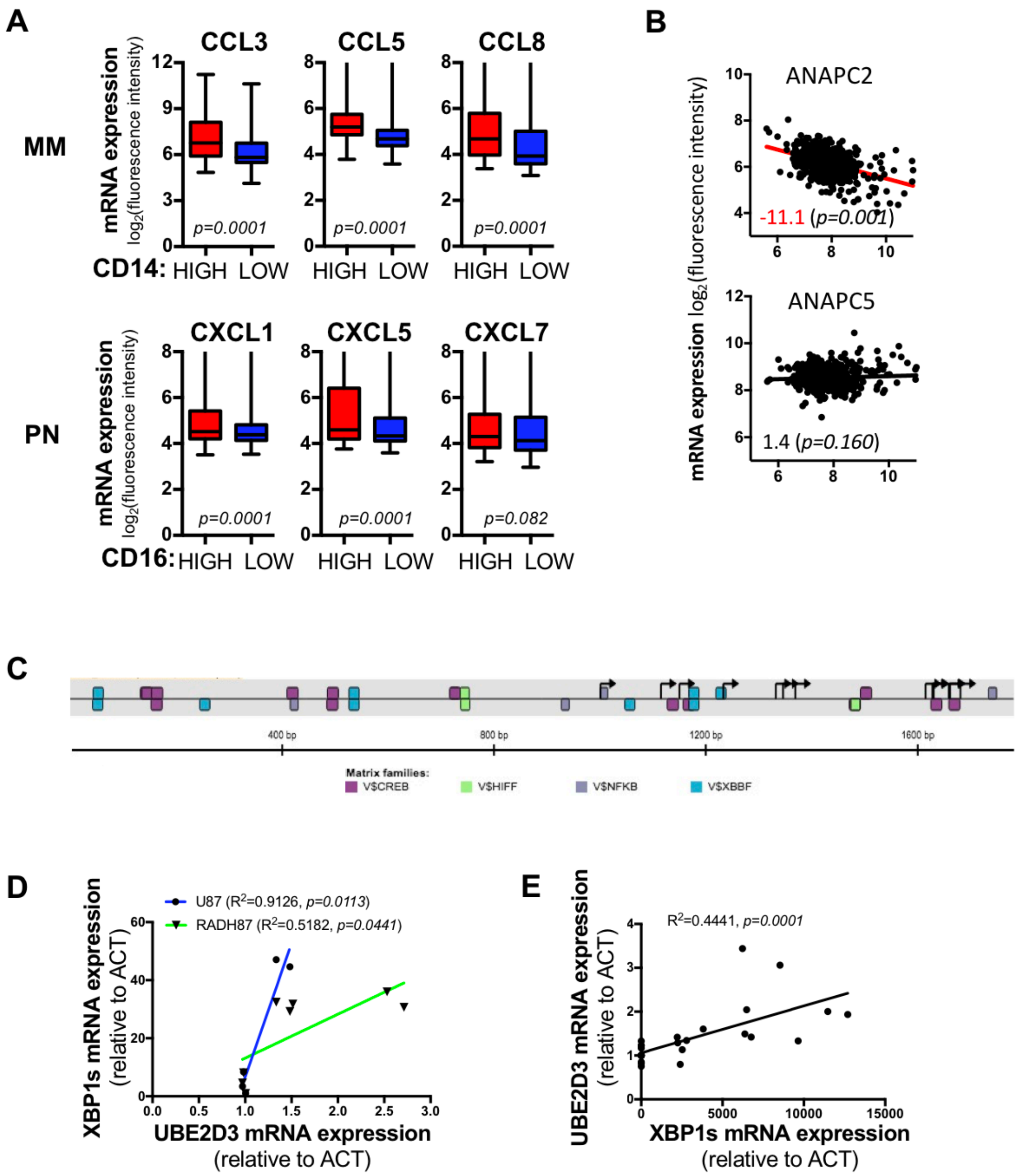

A
MM
PN
mRNA expression
log2(fluorescence intensity)
CCL3
CCL5
CCL8
p=0.0001
p=0.0001
p=0.0001
CD14: HIGH LOW HIGH LOW HIGH LOW
CXCL1
CXCL5
CXCL7
p=0.0001
p=0.0001
p=0.082
CD16: HIGH LOW HIGH LOW HIGH LOW
B
mRNA expression log2(fluorescence intensity)
ANAPC2
-11.1 (p=0.001)
ANAPC5
1.4 (p=0.160)
C
400 bp
800 bp
1200 bp
1600 bp
Matrix families:
V$CREB
V$HIFF
V$NFKB
V$XBBF
D
U87 (R2=0.9126, p=0.0113)
RADH87 (R2=0.5182, p=0.0441)
XBP1s mRNA expression
(relative to ACT)
UBE2D3 mRNA expression
(relative to ACT)
E
R2=0.4441, p=0.0001
UBE2D3 mRNA expression
(relative to ACT)
XBP1s mRNA expression
(relative to ACT)

### Figure S3

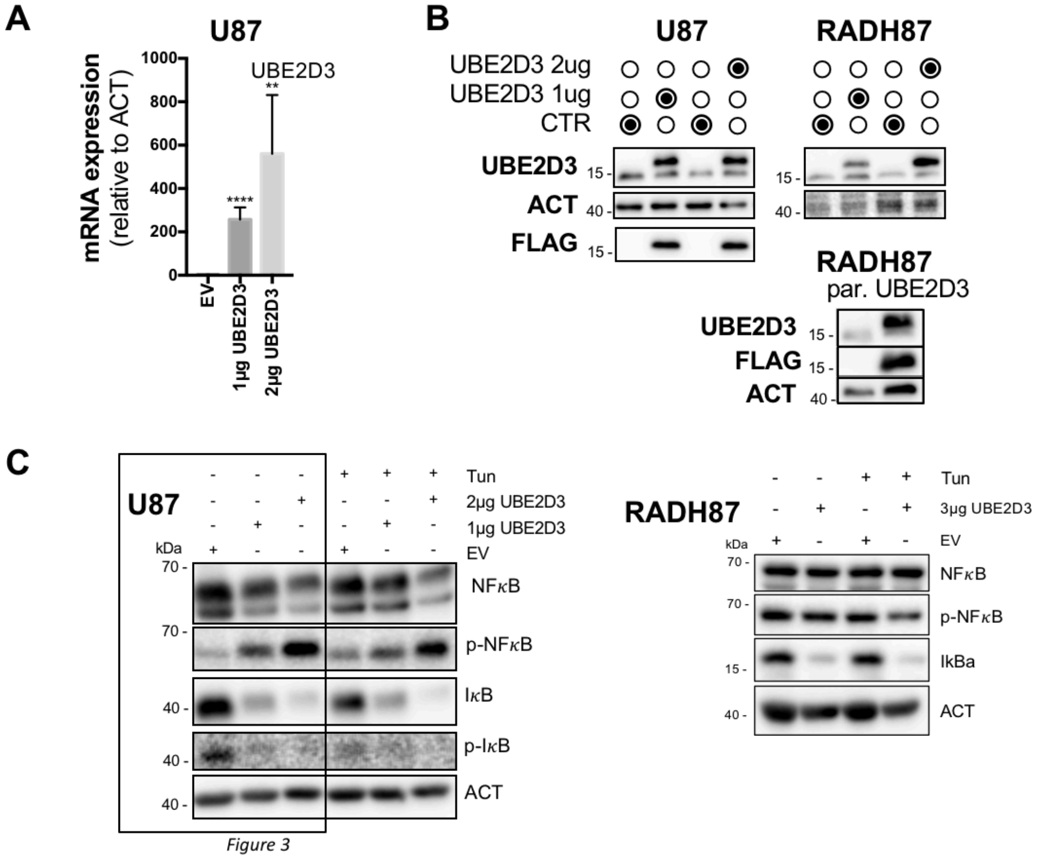
A
U87
UBE2D3
mRNA expression (relative to ACT)
EV
1µg UBE2D3
2µg UBE2D3
B
U87
RADH87
UBE2D3 2ug
UBE2D3 1ug
CTR
UBE2D3
ACT
FLAG
RADH87
par. UBE2D3
C
U87
Tun
2µg UBE2D3
1µg UBE2D3
EV
kDa
NFκB
p-NFκB
IκB
p-IκB
ACT
Figure 3
RADH87
3µg UBE2D3
IkBa

### Figure S4

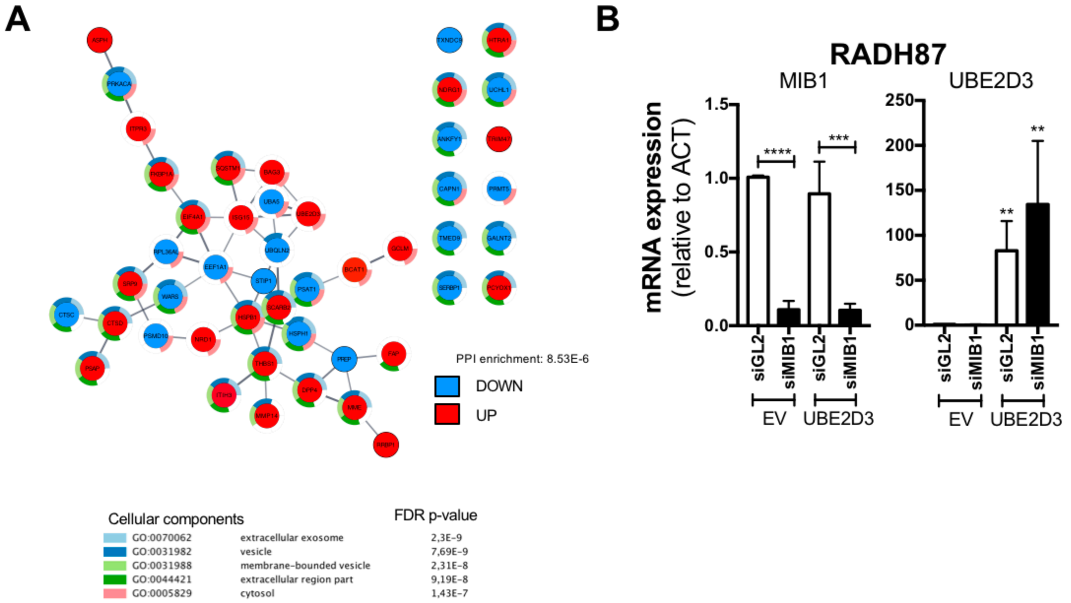
A
PPI enrichment: 8.53E-6
DOWN
UP
Cellular components
FDR p-value
GO:0070062 extracellular exosome 2,3E-9
GO:0031982 vesicle 7,69E-9
GO:0031988 membrane-bounded vesicle 2,31E-8
GO:0044421 extracellular region part 9,19E-8
GO:0005829 cytosol 1,43E-7
B
RADH87
MIB1
UBE2D3
mRNA expression
(relative to ACT)
1.5
1.0
0.5
0.0
250
200
150
100
50
0
****
***
**
**
siGL2
siMIB1
siGL2
siMIB1
EV
UBE2D3

### Figure S5

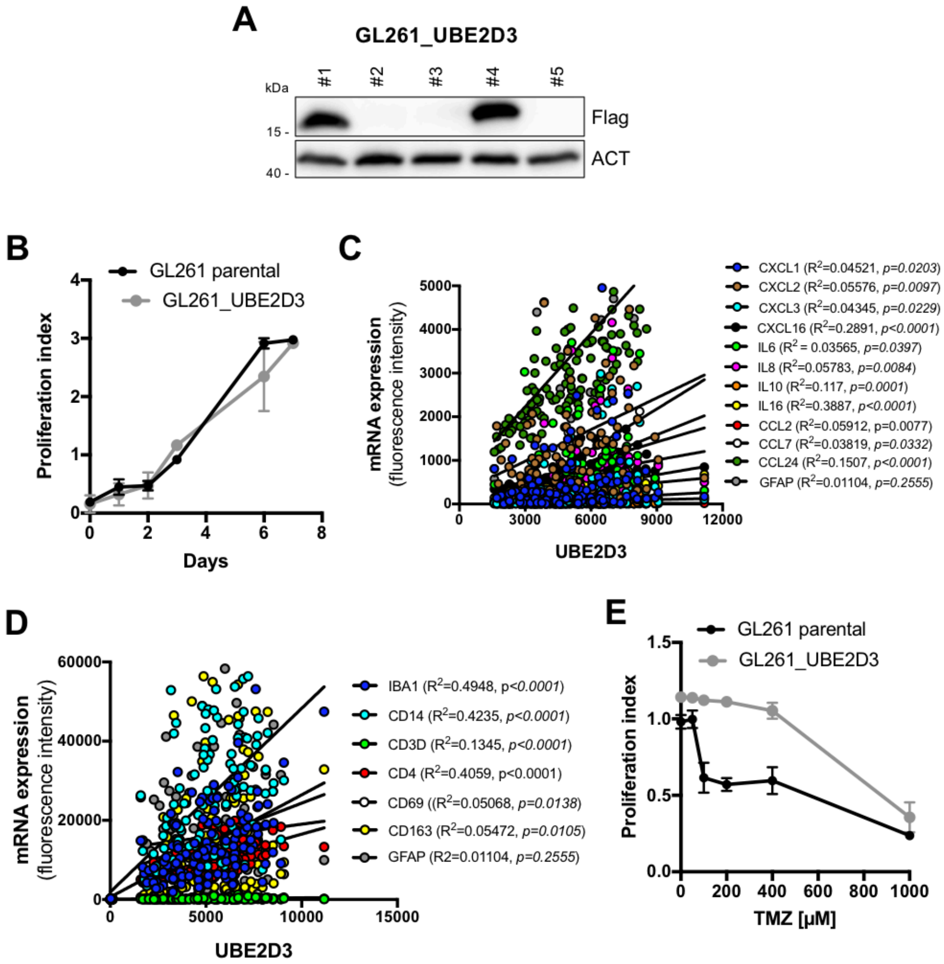

A
GL261_UBE2D3
kDa
#1
#2
#3
#4
#5
15 -
40 -
Flag
ACT
B
GL261 parental
GL261_UBE2D3
Proliferation index
Days
C
mRNA expression (fluorescence intensity)
UBE2D3
CXCL1 (R2=0.04521, p=0.0203)
CXCL2 (R2=0.05576, p=0.0097)
CXCL3 (R2=0.04345, p=0.0229)
CXCL16 (R2=0.2891, p<0.0001)
IL6 (R2 = 0.03565, p=0.0397)
IL8 (R2=0.05783, p=0.0084)
IL10 (R2=0.117, p=0.0001)
IL16 (R2=0.3887, p<0.0001)
CCL2 (R2=0.05912, p=0.0077)
CCL7 (R2=0.03819, p=0.0332)
CCL24 (R2=0.1507, p<0.0001)
GFAP (R2=0.01104, p=0.2555)
D
mRNA expression (fluorescence intensity)
UBE2D3
IBA1 (R2=0.4948, p<0.0001)
CD14 (R2=0.4235, p<0.0001)
CD3D (R2=0.1345, p<0.0001)
CD4 (R2=0.4059, p<0.0001)
CD69 ((R2=0.05068, p=0.0138)
CD163 (R2=0.05472, p=0.0105)
GFAP (R2=0.01104, p=0.2555)
E
GL261 parental
GL261_UBE2D3
Proliferation index
TMZ [μM]
